# Strategies for quantitative RNA-seq analyses among closely related species

**DOI:** 10.1101/297408

**Authors:** Swati Parekh, Beate Vieth, Christoph Ziegenhain, Wolfgang Enard, Ines Hellmann

## Abstract

With the growing appreciation for the role of regulatory differences in evolution, researchers need to reliably quantify expression levels within and among species. However, for non-model organisms genome assemblies and annotations are often not available or have inferior quality, biasing the inference of expression changes to an unknown extent. Here, we explore the possibility to map RNA-seq reads from diverged species to one high quality reference genome. As test case, we used a small primate phylogeny ranging from Human to Marmoset spanning 12% nucleotide divergence. To distinguish the effect of sequence divergence and genome quality, we used *in silico* evolved genomes and existing genomes to simulate RNA-seq reads. These were then mapped to the genome of origin (self-mapping) as well as to one common reference (cross-mapping) to infer the quantification biases. We find that the bias due to cross-mapping is small for the closely related great apes (≤ 4% divergence), and preferable to self-mapping given current genome qualities. For closely related species, cross-mapping provides easy access, high power and a well controlled false discovery rate for both; the analysis of intra-species expression differences as well as the detection of relative differences between species. If divergence increases, so that a substantial fraction of reads exceeds the limits of the mapper used, we find that gene-specific corrections and effect-size cutoffs can limit the bias before self-mapping becomes unavoidable. In summary, for the first time we systematically quantify biases in cross-species RNA-seq studies, providing guidance to best practices for these important evolutionary studies.

## Introduction

Gene expression is an accessible phenotype that can help to bridge the gap between evolutionary changes of the genome and more complex phenotypes [e.g. 1, 2, 3, 4, 5, 6, 7]. Initial genome-wide studies used microarrays to compare closely related species like primates [1, 8, 9] or flies [10] and used genomic information to correct the effect of sequence differences on probe binding [11, 8] or to design species-specific arrays [9, 12]. While microarrays are still a viable approach [6], quantifying expression levels among species by RNA-seq is now preferable. It is expected to provide a less biased quantification of transcriptome-wide expression levels than arrays based on one species and more precise. Additionally, it is less costly than arrays based on species-specific probes.

Nevertheless, systematic technical biases can also occur in RNA-seq studies and hence can bias quantification. For example, the mappability of RNA-seq reads depends on the quality of the genome assembly and the quality of its gene annotation [13]. This could matter, as for example the N50 contig sizes of the current Chimpanzee, Gorilla, Orangutan, Marmoset and Macaque genome assemblies are between 5 and 50 × smaller than the N50 of the current human genome assembly and have three times fewer exons annotated (Table 1).

**Table 1.**
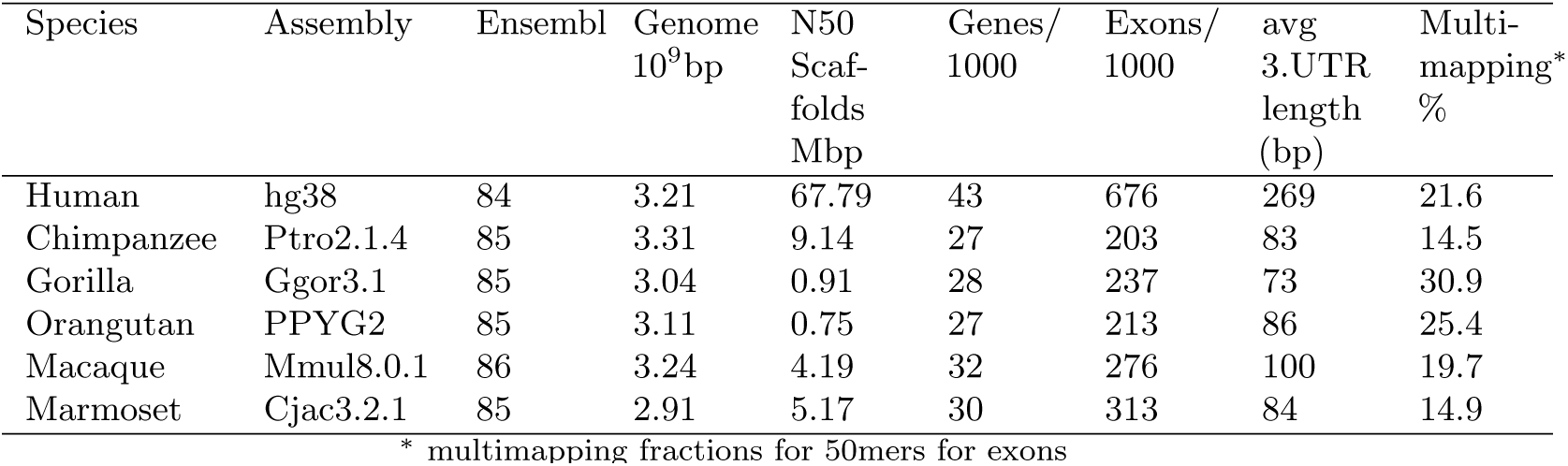
Genome assembly and annotation properties of 6 primates.

An even more difficult part is to define transcriptional units that are comparable across species. Blekhman et al. [14] identified orthologous exons as exons that are present in all three species of that study, i.e. Human, Chimpanzee and Macaque. Going exon by exon rather than attempting to align whole transcripts helped to avoid truncating genes due to bad genome assemblies. In contrast, Brawand et al. [15] opted to find orthologous transcripts. Because transcript annotation is biased due to the use of Human gene models to annotate primate genomes, Brawand et al. [15] used their RNA-seq data to improve the annotation of primate transcripts, before intersecting the transcripts to then only count reads from orthologous parts. This reduction to only common exons or transcript parts also serves as a correction for differing gene lengths between species, however note that this correction is only valid, if reads are evenly sampled across the transcript. Zhu et al. [16] implemented several of the above ideas of identifying and filtering orthologous exons in one pipeline and added an empirical measure of mappability. This approach was used in one of the largest primate transcriptome comparisons to date [17]. Finally, Wunderlich et al. [18] used curated whole genome multiple alignments [19] to convert coordinates of reads mapped to non-human-primate (nh-primate) genomes to Human coordinates and then used the human gene annotation as comparable transcriptional unit.

In summary, efforts so far focused on mapping reads to the genome of each species and counting only reads mapping to a stringently defined set of orthologous exons or genes. What has so far not been investigated is the extend of bias introduced by mapping to different species and to what extend it is possible and even preferable to map to a reference genome from a different species that has a better quality and annotation. Here, we use simulated RNA-seq data from actual primate genomes and *in silico* genomes, that we created by simulating whole genome evolution on a primate phylogeny. The resulting false positive and false negative rates provide recommendations for sequencing and mapping strategies and we can show that mapping RNA-seq reads to one common high quality reference is a viable and sometimes even preferable option for closely related species.

## Results

### Real and simulated primate genomes

We downloaded the Human, Chimpanzee, Gorilla, Orangutan, Macaque and Marmoset genomes from Ensembl [20]. Their average nucleotide divergences to human are 1.3%, 1.8%, 3,7%, 7% and 12.3%, respectively (Figure 1) and their assembly qualities differ considerably with N50 scaffolds ranging from 0.75 Mbp (orangutan) to 68 Mbp (human) and gene annotations range from 676,000 exons and 43,000 genes (human) to 203,000 exons and 27,000 genes (chimpanzee) (Table 1). Another, for our purposes more direct measure of genome quality, is our ability to unambiguously map reads derived from exons. Assuming that the repeat content and thus the uniqueness of kmers is *a priori* similar among primates, we expect most differences in mappability to be due to ambiguous characters in the genome.

**Figure 1.**
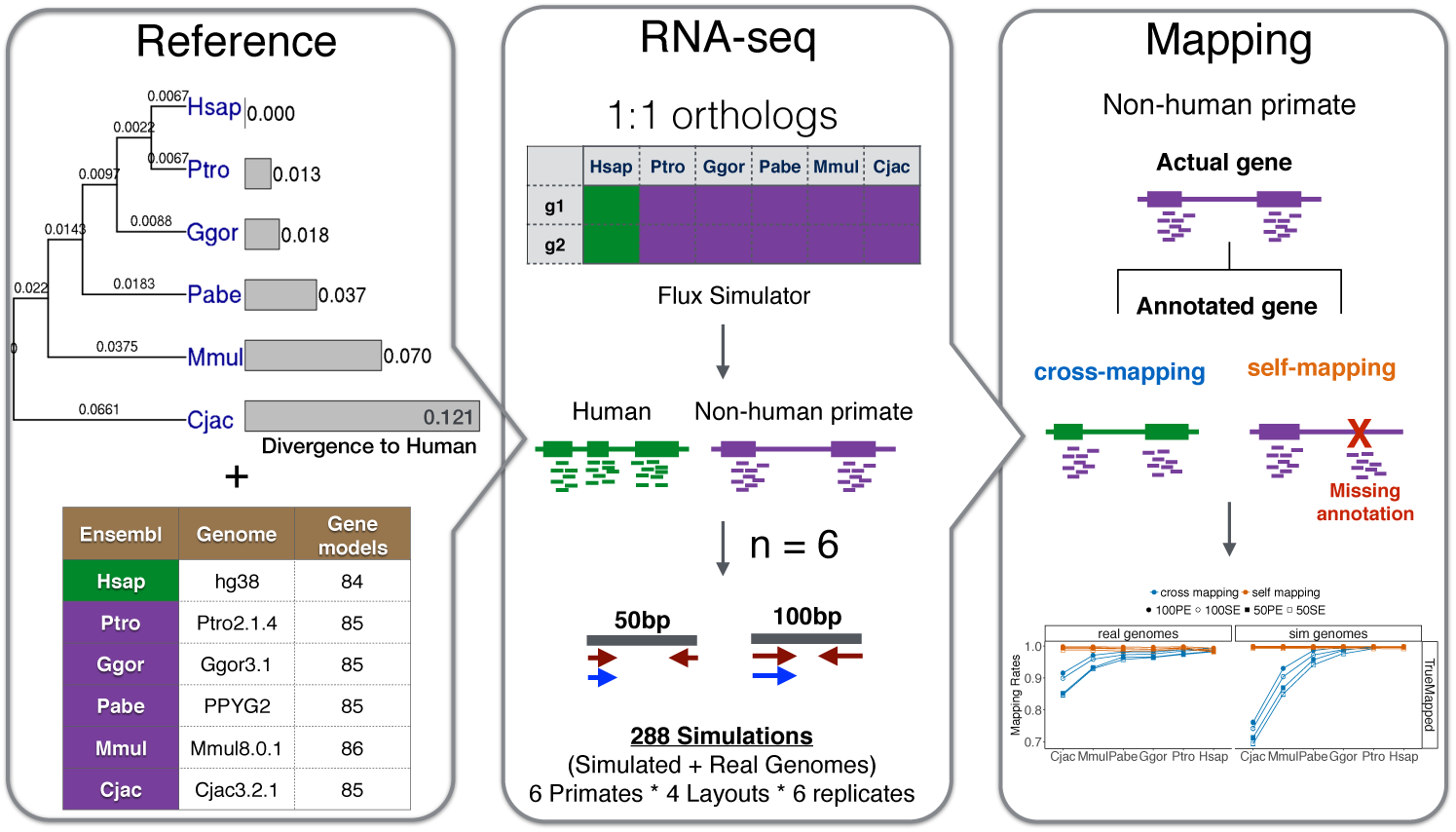
Schematic overview of the study design. **Left)** We use two sets of genome references in this study 1) evolver genomes: *in-silico* evolved genome sequence evolution simulation using hg19 as a common ancestor to simulate 6 primate genomes based on the phylogenetic tree in the left panel: we adopted the nameing from Ensembl with Cjac = Marmoset, Mmul = rhesus Macaque, Pabe = Orangutan, Ggor = Gorilla, Ptro = Chimpanzee, Hsap = Human. 2) The table lists assembly and annotation versions of genomes and gene models downloaded from Ensembl. **Middle)** Based on the *in silico* and Ensembl genomes, we simulate RNA-seq reads using Flux Simulator to generate RNA-seq reads for orthologous genes across primates. Reads are simulated with 6 replicates and 4 sequencing layouts totalling to 288 RNA-seq datasets. **Right)** Mapping strategies 1) cross-mapping: primate reads are mapped to the human reference and 2) self-mapping: reads are mapped to each species’ reference genome. The schematic here shows the under representation of read counts in a gene with missing annotation in non-human primates. “Actual gene” refers to a biologically present gene in the species and “Annotated gene” refers to available information annotated in a GTF (gene models) file in the databases.

When assessing the mappability of exonic 50mers allowing for up to two mismatches [21], we find that 85% of human exons are unique mapping targets, but only 69-85% in the other genomes (Table 1). Using 100mers improves the mappability by on average ∼11%, but mappability remains variable among genomes to an extend that it may bias the quantification of gene expression (Supplementary Figure S1).

One possibility to circumvent problems due to genome quality and annotation, would be to map reads of all species within a study to the one species with the best reference genome. If possible such an approach would also be very valuable when species without a genome or transcriptome assembly are to be included in the study. However, sequence divergence between species is bound to interfere with mapping and quantification of RNA-seq reads, and in reality genome quality and divergence will be convoluted. In order to clearly separate those effects, we decided to simulate whole genome evolution to create realistic *in silico* genomes, using the software evolver [22]. Evolver integrates current knowledge about genome evolution: One can provide rates for various events starting from mutations affecting single nucleotides to large chromosomal rearrangements that can produce duplications of entire genes. The probability of accepting a mutational event is then determined as the inverse of the amount of constraint on the affected sites. Thus mutations at synonymous sites or within 3’UTRs are more likely to remain than mutations that change amino acids or lead to the loss of an entire exon. While events that would cause the complete loss of a gene are not allowed, all other events can occur. We used the Human genome – the most complete primate genome with the most comprehensive annotation – as a starting point (root) and simulated six derived genomes, corresponding to the primate phylogeny given in Figure 1. The simulated genomes allow us to track genes throughout the tree, so that we know the orthologs and do not need to infer them, which is a tricky task [23]. Furthermore, the quality of all simulated genomes is identical, allowing us to assess the effect of sequence divergence alone.

### Mapping to a diverged genome

First, we evaluated which sequencing strategy would be most useful when mapping to a diverged genome. To generate *in silico* RNA-seq reads from the simulated and real primate genomes, we used the Flux Simulator[24]. Flux simulator models all steps of common RNA-seq experiments starting from reverse transcription and ending with the integration of sequencing errors to provide realistically simulated RNA-seq reads in fastq format (Figure S7). We based the parameters that we used for the Flux simulator on a full-length cDNA method (Smart-seq2) with 50bp or 100bp long reads from one end (single-end, SE) or both ends (paired-end, PE) of the cDNA fragments. As gene annotation for the real genomes, we used Ensembl to identify 9,257 human genes that have one-to-one orthologues in each of the five primate genomes [25]. For the simulated genomes we used ∼ 15,000 genes that are known orthologues (Figure 1). As a mapper we used STAR [26] which is currently the best mapper to quantify expression levels by RNA-seq [27].

Evaluating the different sequencing strategies, we find that longer reads improve mapping to the more diverged genomes. Using 100bp instead of 50bp reduces the unmapped fractions by 1.8% and 4% for Macaque and Marmoset,respectively. Surprisingly, the mapping of PE reads did not improve mapping, on the contrary the fraction of unmapped reads increased. Closer inspection of our simulated data showed that mainly reads from genes with exon gain or loss events were affected. Therefore, we loosened the criteria for counting PE reads following an iterative mapping strategy (see Methods) rather than accepting proper read pairs only. This improved the mapping of PE reads (Figure 2). However, compared to SE reads the iteratively mapped PE reads reduced the unmapped fraction by only 0.5% which is not enough to justify the substantially higher sequencing costs. We therefore focus on 100bp SE reads for the remainder of the study. When mapping reads to the simulated genome from which they were derived (self-mapping), we find as expected that almost all reads (*>*99%) map back to the correct location, showing that other factors than divergence and gene annotation play negligible roles. When mapping primate reads from simulated genomes to the simulated Human genome (cross-mapping), we find that between 0.4% (chimpanzee) and 23.6% (marmoset) of reads can not be mapped (Figure 2). Most of those unmapped reads are likely to exceed the divergence level of 10% which is the maximum that is tolerated by STAR. For the mapped reads, the vast majority of reads was mapped correctly and only *<* 2% of all reads map to the wrong location. At the first glance, it may seem surprising that divergence in the simulated genomes has a larger effect on mapping than for the Ensembl genomes, but this can be explained if the one-to-one orthologues of the Emsembl genomes are enriched for more conserved genes. In any case, the fraction of unmapped reads notably increases in the two most diverged genomes (Macaque 2.3% and Marmoset 8.4%), while mapping the great ape reads to the human reference results in almost no loss due to divergence (Chimpanzee unmapped faction:0.6%, Gorilla 0.9%, Orangutan 1.4%; Figure 2).

**Figure 2.**
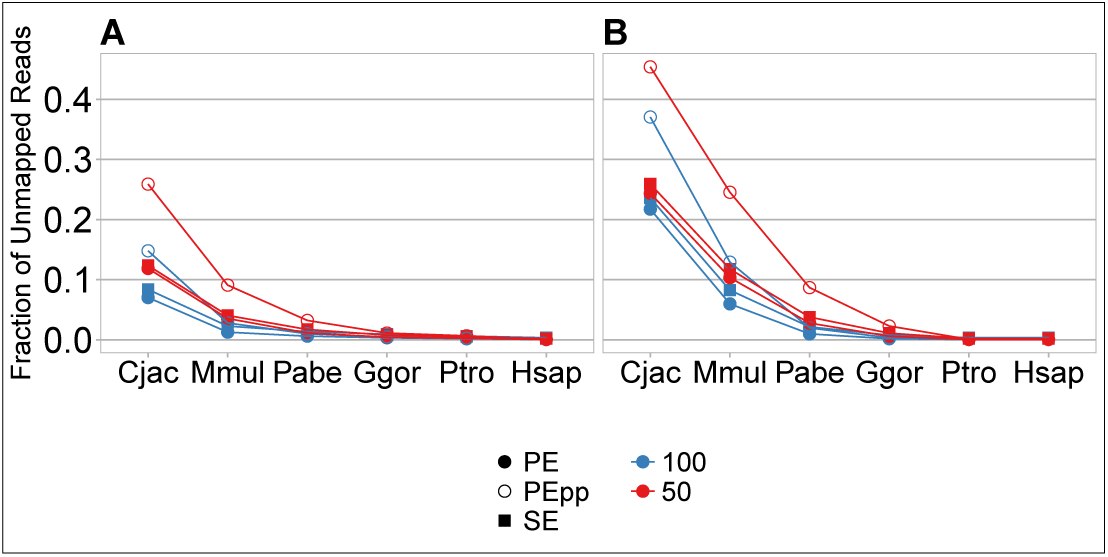
The fraction of unmapped reads for different sequencing strategies. The fraction of reads that remain unmapped when mapping primate reads to the Human genome (cross-mapping) for RNA-seq reads simulated from Ensembl genomes **A** and *in silico* genomes **B**. We also compare the impact of different sequencing strategies on the fraction of unmapped reads of 50bp (red) and 100bp (blue) reads. Squares are single-end and circles are paired-end layouts. Empty circles represent the statistics if only proper pairs were kept, the filled circles represent an iterative approach.

### Impact of gene-wise divergence on mapping

If the number of reads that is lost due to mapping to a diverged genome was evenly distributed across all genes, this would have little impact on measures of differential expression: The other genome would simply appear to have fewer reads in total and standard normalization procedures would take care of such a discrepancy. Problems will only arise, if due to divergence some genes lost more reads than others, i.e. it is not the total divergence that is of interest, but the variance in divergence - and thus mapping success - across genes.

To this end we use the cross-mapping data from the *in silico* genomes to estimate divergence (see Methods) and check whether divergence correlates with the correct mapping fraction. Note that estimating divergence from the RNA-seq data itself makes it possible to obtain divergence estimates for species without other available genome or transcriptome sequences.

Indeed, we find that the fraction of correctly mapped reads has an inverse correlation with gene-wise divergence and this correlation increases with species divergence (Figure 3). Chimpanzee and Human are so closely related that the little divergence has no discernible effect on mapping success (Pearson’s *r_Hsap_* = −0.01*, p* = 0.28), while for Marmoset and Macaque the fraction of correctly mapped reads strongly depends on sequence divergence of the gene (*r_Cjac_* = −0.50∗∗∗, *r_Mmul_* = −0.39∗∗∗).

**Figure 3.**
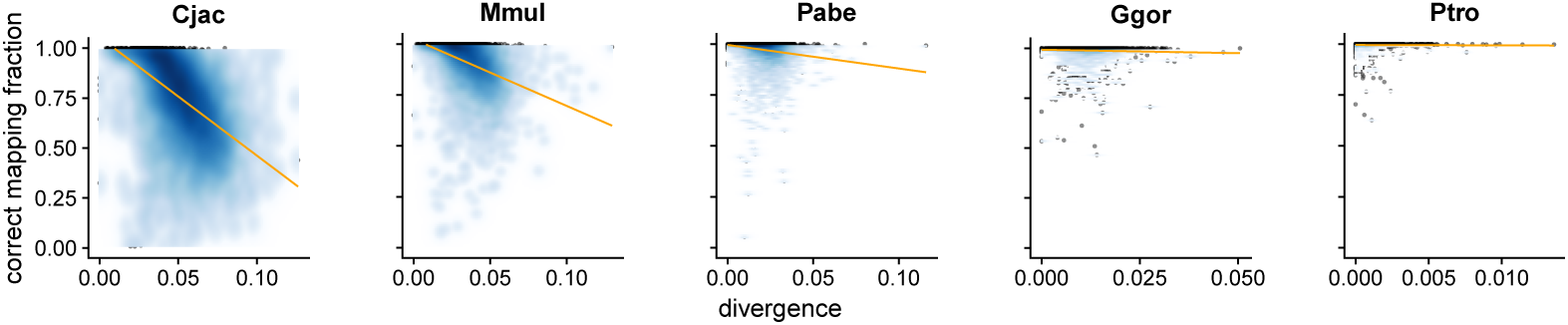
Gene-wise divergence impacts the fraction of correctly mapped reads. Divergence was estimated for genes with *>* 100 simulated reads. The fraction of correctly mapped reads, is the fraction of reads that could be recovered from the ones that were sampled from that gene. The orange line is the linear regression of the divergence on the mapping fraction.

Based on this result, mapping Chimpanzee reads to the human genome is not expected to bias transcript quantification, while there will be an effect for Marmoset and Macaque. However, it might be possible to use the divergence estimates to correct the read counts obtained by cross-mapping.

### Impact of cross-mapping on differential expression analyses

While our analysis shows that cross-mapping quantification of expression levels are likely to be accurate for closely related species, it is unclear how cross-mapping of more diverged species would affect the identification of differentially expressed (DE) genes. To this end we used Flux Simulator [24] to generate 100bp SE RNA-seq reads for orthologous genes from the six *in silico* genomes, keeping the expression levels fixed across all species. We simulated 6 replicates/species and tabulated the number of significantly differentially expressed genes between each of the five primate RNA-seq datasets to Human, using DESeq2 [28]. Note that since we did not simulate any expression changes, all DE-genes detected are false positives.

Reads were mapped to both, the genome of origin (self-mapping) as well as the *in silico* Human genome (cross-mapping). To test whether one can correct for lost reads due to divergence using the RNA-seq data, we used a log-linear model to predict the number of sampled reads from the number of mapped reads per gene given gene-wise divergence estimates (Gene-wise divergence estimation; see Methods).

The number of false positives using self-mapping is negligible (up to 0.1% Marmoset, (Figure 4). This is not surprising since the simulated genomes have no errors and a precise annotation leading to near perfect mappability. Furthermore, unlike for the real Ensembl genomes there is no doubt about orthology. Thus, all false positives detected with cross-mapping must be due to sequence divergence.

**Figure 4.**
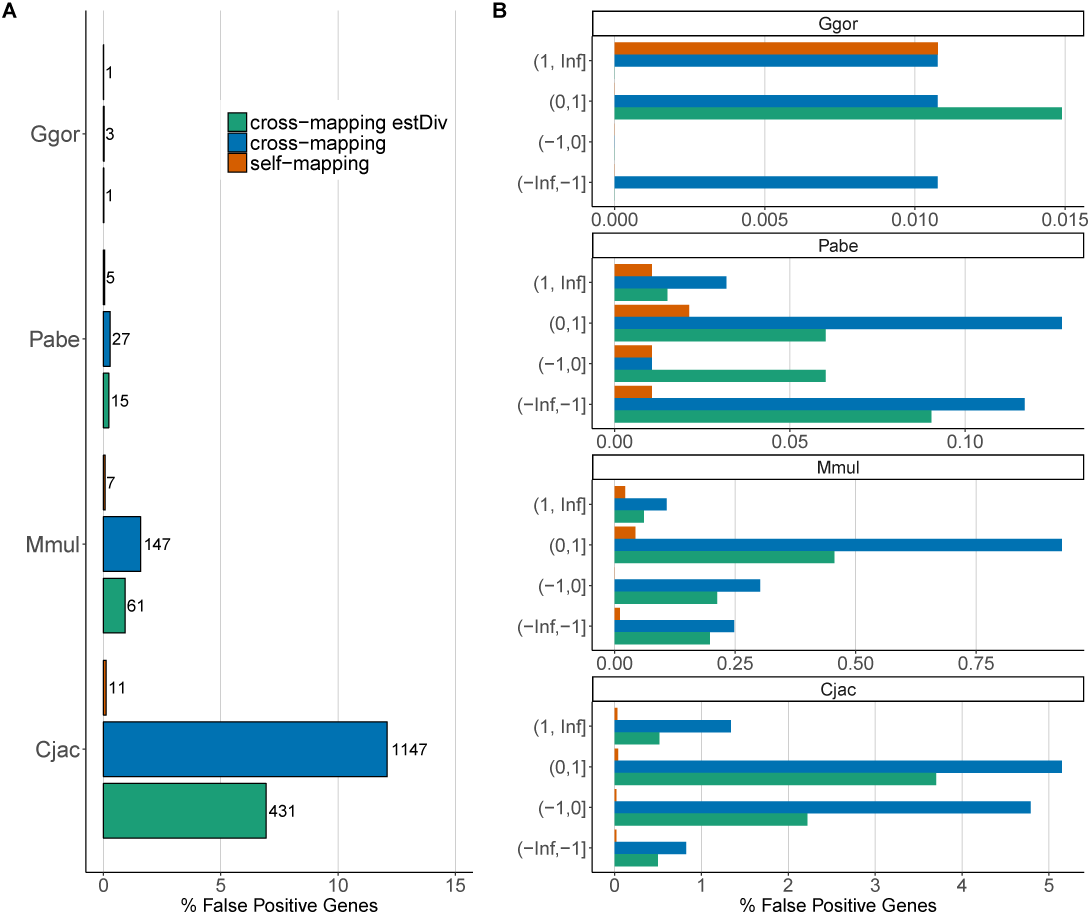
False Positive DE genes for different counting strategies for *in-silico* genomes. We simulate 100bp SE RNA-seq reads from *in silico* evolved genomes. We consider three counting strategies: 1) Reads were either mapped to the genome of origin (self-mapping, orange) 2) reads were mapped to the *in silico* Human genome (cross-mapping, blue) and 3) cross-mapping counts were corrected for divergence estimates (cross-mapping estDiv, green). We used DESeq2 [28] to find DE-genes between Human and the non-human primate (FDR 0.05). Panel **A** shows the marginal FPR for Gorilla-Human (Ggor), Orangutan-Human (Pabe), Macaque-Human (Mmul) and Marmoset-Human (Cjac) comparisons for all three counting strategies, including the numbers of false positives genes for each comparison. For the Human-Human and Chimpanzee-Human comparison, no false positives were found. In panel **B**, we stratify the FPR according to the estimated *log*_2_-fold change. A positive *log*_2_-fold change indicates a higher and a negative a lower expression in the Human (reference) as compared to the non-human primate.

With cross-mapping, the false positive rate (FPR) increases with divergence (Figure 4). While for both Human and Chimpanzee we did not observe any false positives, Gorilla and Orangutan have acceptable FPRs with 0.04% and 0.34%, respectively. This rate increases to 2% and 14% for Macaque and Marmoset for which an increasing number of genes have unmapped reads due to high divergence (Figure 3. This also results in an increasing number of genes that are biased towards higher expression in Human, i.e. in the reference species (Figure 4A, Supplementary figure S2). ∼60% of all false positives appear to have a higher expression in Human than the non-human-primate. Using our gene-wise divergence estimates to correct for variation due to unmapped reads, the FPR is reduced by ∼ half (Macaque 1.2%; Marmoset 7.9%): While clearly an improvement, it does the FPR still increases with species divergence, is high compared to self-mapping and biased towards the reference species (Figure 4A, Supplementary figure S2).

We identified two reasons, why our divergence correction does not work as well as we hoped. On the one hand, we can only estimate divergence from reads that could be mapped (Figure 3), which leads us to underestimate divergence for highly diverged genes and thus an underestimation of the number of sampled reads. On the other hand, the small fraction of falsely mapped reads will lead to an overestimation of divergence and thus an overestimation of sampled reads (Supplementary Figure S3). Indeed, especially the false positive genes that also have a high *log*_2_-fold change, show an enrichment for falsely mapped reads (Supplementary Figure S4)

However, even though divergence to the reference genome introduces significant false positives, the effect size of the majority of those changes is small. Most *log*_2_-fold changes are between 1 and −1, and thus smaller than an absolute 2-fold difference (Figure 4B). If we require genes to have a significant absolute *log*_2_-fold change of at least 1, the FPR for cross-mapping counts reduces to 0.3% in Macaque and to 1% in Marmoset.

In summary, our simulations suggest that mapping to a diverged genome for quantitative expression analysis is clearly a viable option for genomes that are less than 4% diverged. At higher divergences, the FPR increases to more and more substantial levels. Using a model that corrects for the observed divergence and by filtering for larger effect sizes, the FPR can be reduced, but the increasing number of falsely mapped reads limits the amount of correction possible for more diverged genomes.

### DE-analysis across species for Ensembl-genomes

Next, we wanted to investigate in how far we can generalize our findings to the more realistic scenario, with varying genome and annotation quality. To this end we simulate RNA-seq reads from the ‘real’ primate genomes as downloaded from Ensembl (Table 1), otherwise the experimental setup and the DE-analysis were the same as for the *in silico* genomes and should thus be comparable.

To begin with, the self-mapping FPRs are much higher for the Ensembl than for the *in silico* genomes (Ensembl: 1.5-2.6% vs. *in silico ∼*0.1%). Part of this discrepancy might be due to differences in gene lengths that, due to annotation issues, appear exaggerated in the Ensembl genomes. For example, 3’UTRs are consistently shorter in the non-human primates (Table 1). In order to correct for these gene length differences, we only counted reads mapping to orthologous sequences that were annotated in both the Human and the non-human primate [15, e.g.]. This helped to reduce the FPR to ∼1% in all comparisons (Figure 5). When using cross-mapping counts, the FPRs for the great ape genomes are low: With 0.6% (Chimpanzee), 0.7% (Gorilla) and 1% Orangutan, cross-mapping FPRs are even lower than self-mapping counts (Figure 5).

**Figure 5.**
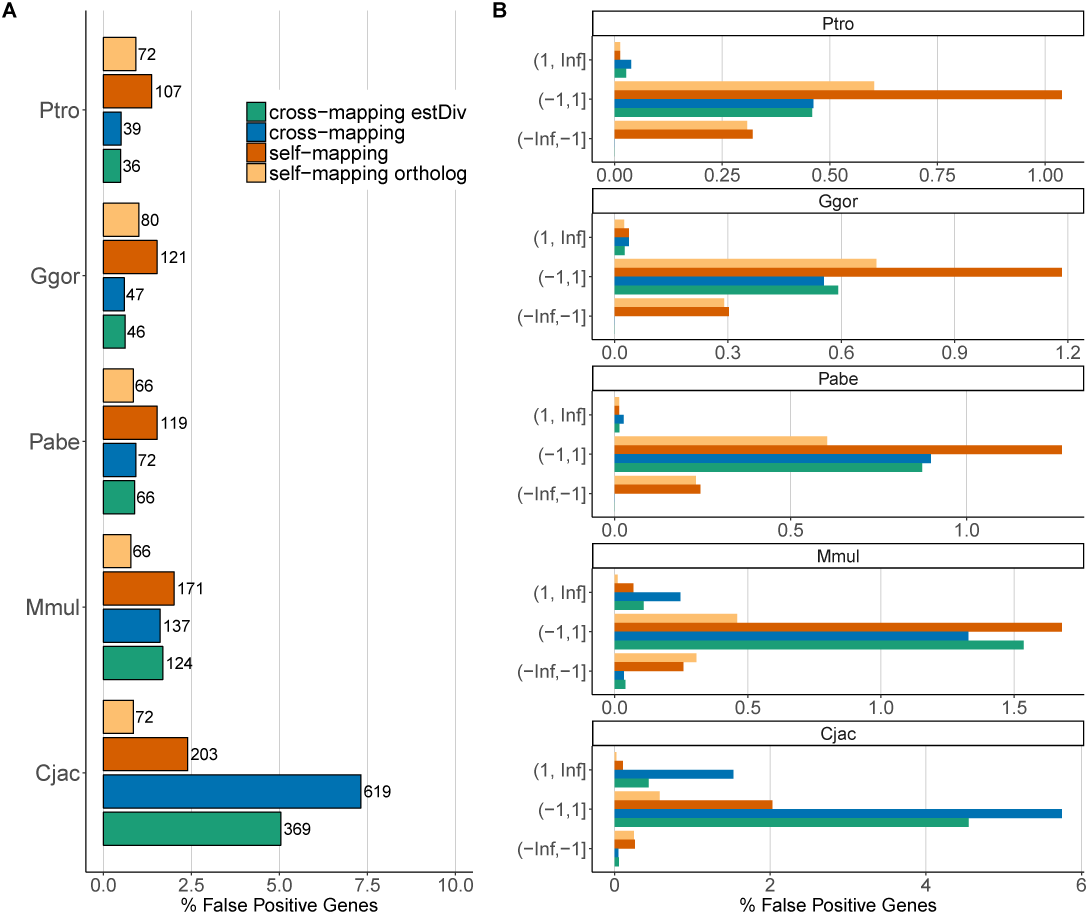
False Positive DE genes for different counting strategies for Ensembl genomes. This plot is the same as Figure 5, except that here we simulate RNA-seq reads from Ensembl genomes (hg38, Ptro2.1, Ggor3.1, Ppyg2, Mmul8.0.1, Cjac3.2). Furthermore, here we investigate two counting strategies for the self-mapping reads: 1) self-mapping simply counting all reads that were mapped to orthologous genes (orange) and 2) self-mapping, counting only reads that map to the orthoglous parts of the genes (self-mapping ortholog, yellow).

As in the simulated genomes, the FPRs for Macaque (2%) and Marmoset (8.5%) are much higher than for the great apes. The FPR for Macaque agrees with the level estimated for the *in silico* genomes, while the FPR from the *in silico* Marmoset genome (14%) is substantially higher. This suggests that the set of one-to-one orthologs is biased towards conserved genes, and the Marmoset as the most diverged species in the comparison is the limiting factor to find one-to-one orthologs across all species. Furthermore, the directional bias in the false positives is the most pronounced for the Marmoset, where 3 times more false positives appear to be up-regulated than down-regulated in the Human (Supplementary Figures S5 & S6). Using divergence estimates to correct cross-mapping counts does not reduce the FPR in Macaque, and provides only a slight improvement by 2.2% for Marmoset (Figure 5), suggesting that in practice a simple divergence correction without further mappability assessment will be insufficient.

Finally as for the *in silico* genomes, the effect sizes of the false positives are also low for the Ensembl genomes and instating a cut-off for absolute *log*_2_-fold changes of at least 1 reduces the FPR for all cross-species comparisons to ≤ 0.5%.

### Within species DE-analysis while mapping to a diverged reference

Although divergence somewhat impairs DE-analysis between species, it is unclear how mapping to a diverged genome would impact DE-analysis within a species. This would allow to conduct DE-analysis for species for which only a genome of a close relative has been assembled. To test such an analysis strategy, we simulate RNA-seq data for each of the six simulated genomes and two conditions for each species. 10% of the genes are simulated to have a 4-fold change between conditions, where equal numbers of genes are up and down regulated (Figure 6). Again we map all reads to the Human genome and analyze differential expression using DESeq2 to tabulate how many of the simulated DE-genes can be recovered (=sensitivity, true positive rate, TPR) as well as the proportion of false discoveries among all significant genes (=specificity, false discovery rate, FDR). The mapping of human reads to the human genome yielded a TPR of 99% and an FDR of 2.5%. This represents the best case scenario, i.e. self-mapping to a high quality, well annotated genome. Chimpanzee reads yield similarly good sensitivity and specificity. Mapping Gorilla, Orangutan and Macaque reads to the Human genome also yields a TPR close to 99%, but the FDR appears slightly increased to 3% in Gorilla and Orangutan and 4.5% for Macaque. However, only Marmoset with an FDR of 6% exceeds the nominal level and also shows a noticeable drop in TPR to 97%. Because both conditions are affected by divergence to the same extend, divergence correction does not help to reduce the FDR but only decreases sensitivity (Figure 6).

**Figure 6.**
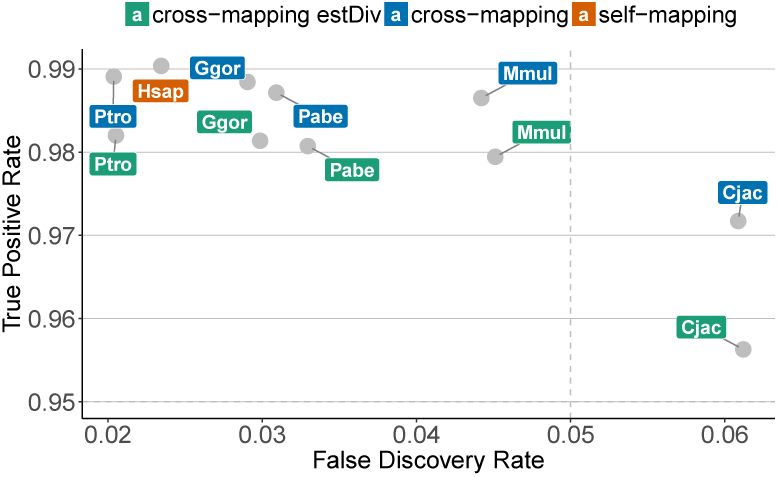
Within-species DE-analysis mapping to a diverged Genome. We simulate reads for 2 conditions with 10% DE-genes using the *in silico* genomes and assess the true positive rate (TPR) and the false discovery rate (FDR), for our three counting strategies: self-mapping (yellow), cross-mapping (blue) and divergence corrected cross-mapping (green).

In the absence of of a good reference genome, mapping to a diverged reference for within species DE-analysis yields good sensitivity and only the FDR for species as diverged as Humans and Marmosets starts to be problematic.

### Detecting relative expression changes between species

In many instances, changes in the regulation of a gene are easier to interpret than absolute expression differences [29]. For example, if one gene is up-regulated during development in one species, but not in the other, the first time-point serves as an internal reference that facilitates the detection of this regulatory difference. This makes the inference of an inter-species difference more robust towards technical biases, including systematic species differences.

For differential expression analyses tools based on (generalized) linear models [28, 30] such a regulatory difference would be formulated as the interaction term between species and condition, i.e. the expression change between conditions depends on the identity of the species. In the previous chapter, we described the simulation of two conditions within each species. Note that we simulated no expression changes between species, in other words condition A, as condition B, should have identical expression profiles for both species (Figure 7). Hence, also the expression changes between conditions occur in both species and thus a significant interaction term species:condition represents a false positive regulatory change.

**Figure 7.**
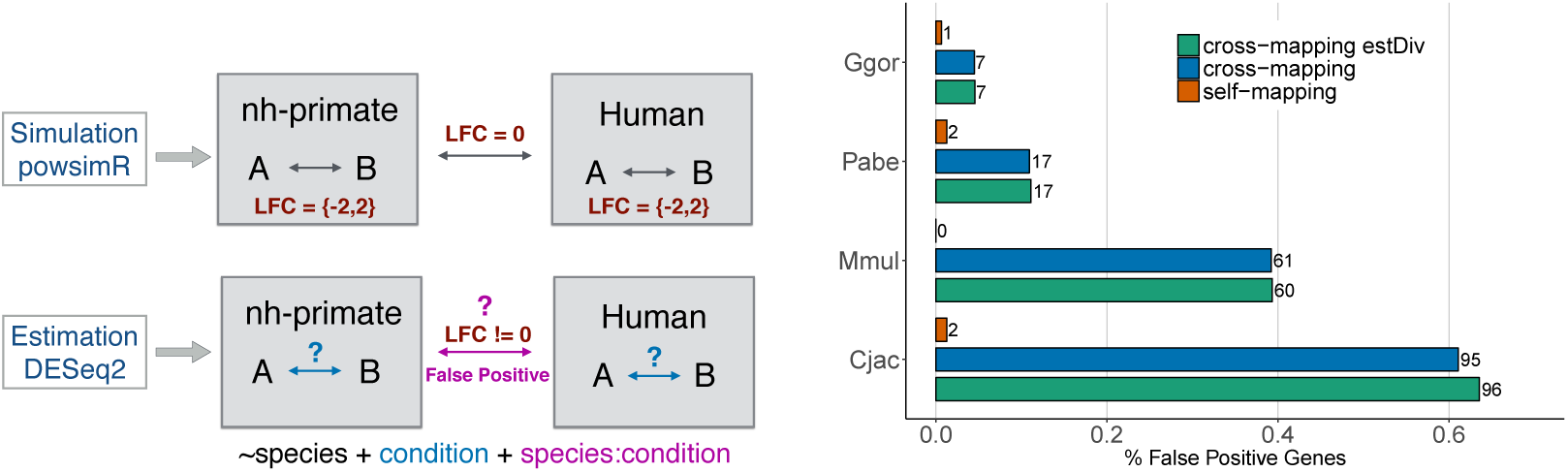
Relative expression measures. **A)**The upper panel shows simulation and the lower panel estimation schematic. We simulate RNA-seq reads for each non-human primate - Human pair for two conditions, whereas 10% of the genes have a 4-fold expression difference between conditions. There should be no differential expression between species. We then used DESeq2 to find DE-genes between conditions in each primate and condition difference between species. **B)** The barplot shows the false positively inferred relative expression changes between species, i.e. we detect a gene to be DE between conditions in one species but not the other. The colors indicate counting strategies (self-mapping - Orange, cross-mapping - blue and cross-mapping with divergence correction - green).

Again in the comparison between Human and Chimpanzee, there are no false positives, and the FPRs for the other great apes with 0.05% in Gorilla (7 genes) and 0.12% in Orangutan (19 genes) are also very low. Even though Macaque (FPR=0.4%) and Marmoset (FPR=0.6) have higher FPRs than the great apes, they are still low compared to the FPRs for the comparison of absolute expression changes (Figure 4A). Furthermore, divergence correction had no impact on the FPR, confirming that a relative model efficiently corrects for species differences that are due to divergence.

Hence, even though FPRs increase with divergence also for relative expression changes, it is much better controlled than for absolute expression changes. Thus keeping an cautious eye on the FPR levels, also comparisons for more distantly related species such as Human and Marmoset will yield meaningful results.

## Discussion

RNA-seq is a versatile tool that also has many applications in evolution and ecology [31]. However, for many of the organisms of interest, no reference genome exists and even if a reference genome is available, the quality of the genome and its annotation can be highly variable. One simple solution to this problem would be to use an available high quality reference genome from a closely related species. Here, we investigate the biases that come by mapping RNA-seq reads to a diverged genome for three types of biological problems: 1) Finding expression differences between species, 2) Finding expression differences within the non-reference species and 3) Finding relative expression changes between species.

We illustrate our approach using a small primate phylogeny including Chimpanzee, Gorilla, Orangutan, Macaque, Marmoset and Human, with neutral divergence ranging from 1.34 −12.1% (Figure 1). In order to distinguish between problems due to annotation and actual evolutionary differences, we simulate RNA-seq data from both primate Ensemble as well as *in silico* evoloved genomes and map them to their Human counterpart (cross-mapping). The *in silico* genomes should only show sequence differences due to ‘evolutionary’ divergence, but not due to assembly or annotation errors. Furthermore, ortholgous genes in the *in silico* genomes are known and hence do not show a bias to higher conservation as we saw in the inferred orthologues in the real genomes.

Sequencing of longer reads improves cross-mapping, but due to possible changes in the gene model, like loosing an exon, paired-end sequencing requires a non-standard mapping strategy and also does not improve the correct mapping fraction enough to justify the higher costs. However, irrespective of the mapping strategy, the fraction of reads that can not be mapped increases with divergence (Figure 2).

That being said, a genome-wide reduction in the number of mappable reads does not necessarily introduce bias into expression analyses. Bias is only introduced, if the fraction of correctly mapped reads varies systematically among genes and we can indeed show that there is an inverse relationship between the divergence of a gene and the fraction of correctly mapped reads (Figure 3). Thus, using cross-mapping counts, we find that more diverged genes appear to have a lower expression in primates as compared to Humans (Figure 4, Supplementary Figure 2). To correct for this effect, we try to regress divergence out of the read count estimates, using gene-wise divergence estimates from the same RNA-seq data. The FPR for these divergence corrected read-counts was roughly half of what it was without divergence correction.

However, especially for the more diverged species (Marmoset and Macaque), some bias towards higher counts in Humans remains. We believe that the reason for the remaining false positives and their directional bias is an underestimation of the true divergence of a gene. Both Marmoset and Macaque have genes with divergence of *>*10%, which marks a threshold from which on many short read mappers including STAR begin to have problems to place reads correctly [26]. Because we cannot map to regions with high divergence, those cannot be included in the divergence estimation, thus making our correction insufficient. Furthermore, bad divergence estimates can also explain some of the additional false positives with much higher counts in primates. This is because mis-mapped reads will inflate our divergence estimates and trigger the regression model to over-correct the read-count (Supplementary Figure S3 & S4).

Fortunately, false mapping rates are low, so that only few genes will show this rather counter-intuitive pattern. Generally, false positive DE-genes have low *log*_2_-fold changes, so that imposing a cut-off on the effect size gets rid of the vast majority of false positives, even for cross-mapping of Marmoset reads to Human (Figure 4).

Most of the results obtained using *in silico* genomes could also be recapitulated with the Ensembl genomes. The main difference, that we saw was a substantially higher FPRs for self-mapping in the real genomes. This discrepancy is probably due to an exaggeration of the differences in the gene models due to annotation problems. Restricting the regions to only the ones that are also annotated in the primate genome should correct for the resulting gene-length differences (Figure 5). However, this correction assumes that RNA-seq reads are evenly sampled across the entire transcript length, which in practise is rarely the case [32]. Thus, for very closely related species such as Chimpanzee and Gorilla, cross-mapping actually produces fewer false positives than self-mapping. For very closely related species the benefits of a good annotation and genome quality outweigh the problems introduced by sequence divergence. In summary with respect to the first biological problem, we can say that cross-mapping can be used to find between species expression differences, but specificity decreases with divergence to the reference.

The second question that we investigated was in how far cross-mapping can be used to analyze differential expression within the same species. To this end we simulated RNA-seq reads for two conditions that differ in their expression profiles and map the reads to a diverged reference. This strategy yields a high sensitivity of over 97% for all species and only the Marmoset had an FDR slightly above the nominal level, showing that the cross-mapping bias cancels out when contrasting two conditions within the same species.

Following up on this notion, we investigated in how far relative differences between species can be detected using cross-mapping. To this end, we simulated RNA-seq data for two conditions in two species, where the conditions do not differ between the species, i.e. also all regulatory changes between conditions are the same in both species and thus we expect no relative changes (Figure 7). Even though the FPR for relative expression changes between species increases with species divergence, it is ∼ 10× lower than the FPR for absolute between species changes.

Hence, in the absence of a good genome assembly or annotation cross-mapping to a closely related species is a good alternative to analyze expression differences between conditions within the same species or to detect relative differences between species. As a general rule of thumb, cross-mapping works well as long as the divergence does not exceed the limits of the mapper used. If divergence for all genes remains below this threshold, no further corrections are necessary. However, if some genes exceed the divergence that can be safely handled by the mapper, quantitative comparison between species will produce more false positives, which can be dealt with by introducing an effect size cut-off.

## Methods

### Generating *in silico* Genomes

We used evolver [22] to simulate whole genome sequence evolution. Given an ancestral genome, annotation of gene models, CpG islands, a repeat library and rates for 1) nucleotide and 2) amino acid substitution, 3) indels, 4) tandem repeat expansion and 5)contraction, evolver models sequence evolution of an entire genome. To simulate gene expression of diverged species using RNA-seq we explicitly model the evolution of coding regions, UTRs, start and stop codons as well as splice junctions. All possible types of mutations can affect functional regions, but evolver constrains their number based on a set amount of selection on the affected sites. Hence genes can also be rearranged or duplicated and exons can be gained or lost.

To evaluate the effect of mapping and quantification for diverged species on realistic data, we used human genome (hg19) and gene models (GRCh37.75) as ancestral genome. Starting with hg19 at root, we generated six *in-silico* genomes based on the neutral divergence estimates for a primate phylogeny (Paten et al. [19],Figure 1).

We used the hg19 parameter file provided by evolver to set up all rates to model evolution and followed the steps described in the evolver manual to prepare the input for the simulations. First, we replaced all ambiguous bases in hg19 by random bases. Next, we obtained gene models in gff format from the UCSC genome browser [33] and filtered overlapping genes, leaving us with non-overlapping 15,559 genes. Because evolver only goes along one trajectory, we simulated evolution successively for each branch, starting with the root and using default parameters. All simulated genomes are converted to fasta format and the evolved gene-models are output in gff format, which are then used for RNA-seq simulation.

### RNA-seq simulation

RNA-seq reads for 6 replicates for all 6 primates were simulated using Flux Simulator [24] from *in-silico* as well as the Ensembl genomes [34] (Figure 1). Reads were generated in single-end and paired-end format with read lengths of 50 and 100 bp each. The error models used in Flux Simulator were built from bulk Smart-seq2 data sequenced on Illumina HiSeq1500 (Supplementary figure S7). In order to simulate equal numbers of expressed mRNA molecules for each orthologous gene across species, we first simulate 10 Million reads for one sample using the zipf-distribution as implemented in Flux Simulator and then use these read counts as input for all samples.

In order to simulate RNA-seq reads for two different conditions, we used molecule counts from 12 bulk UHRR samples from previously published dataset [32] to estimate mean and dispersion relation to simulate relasitc expression distributions using powsimR [35]. The two groups differ in 10% of the genes with equal proportions of 4-fold up- and down-regulation. These molecule counts are then used as input for Flux Simulator to simulate RNA-seq reads. In this simulation framework, we there are again no expression difference between species while expression within species differs between conditions.

### Mapping and Quantification

The simulated RNA-seq reads were mapped using STAR 2.5.3a [26] in “–twopassMode Basic” mode with default parameters. For paired-end layout, we use an “iterative mapping” strategy. First, the reads were mapped as paired-end and unmapped reads were extracted and mapped again as single-end reads, ignoring the information of their mate. We employed two different mapping strategies: The one being self-mapping where reads from different primates were mapped to their own genomes and gene models. The other being cross-mapping primate reads were mapped to the Human genome using Human gene models. In order to correct for annotated gene length differences in the Ensembl genomes, read counting was restricted only to the orthologous regions annotated in human and non-human primates. The orthologous regions identified using reciprocal liftOver [36] between human and non-human primate genomes.

The mapped reads were assigned to genes using featureCounts from the bioconductor package Rsubread [37].

### Gene-wise Divergence estimation and correction

We estimate divergence of each non-human primate gene to its human ortholouge from reads cross-mapped to human. To begin with we combined the alignments for all the replicates of one species and then followed the GATK best practices [38] for SNP-calling from RNA-seq data to improve alignment quality. BAM files were prepared using “AddOrReplaceReadGroups” and “SplitNCigarReads” and then put through GATK “IndelRealigner” with default parameters. From those BAM files, we then created vcf files including invariant sites “samtools.1.3.1 mpileup −Q 0 −uv”. We use a custom R script to estimate gene wise divergence from the above VCF files, wherein we also consider genotype qualities to correct for call-ability.

We then used the divergence estimates to adjust the abundance level of genes for sequence divergence to the reference. To this end, for each gene (*i*) we fit a linear model to predict the simulated counts from Flux Simulator (*y*_*i*_) using the observed cross-mapping counts (*x*_*i*_) and the divergence estimate (*d*_*i*_) (Equation 1).

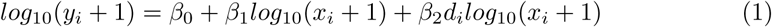

Substitutions are only counted for bi-allelic sites. To obtain divergence estimates, we us Kimuras two parameter model to correct for multiple hits [39].

### Differential Gene Expression

We performed pairwise differential expression analyses of each non-human primate to human from cross mapped counts on simulated data using DESeq2[28]. We used false discovery correction according to Benjamini Yekutieli [40] and a nominal α = 0.05. We performed the differential expression analysis for (i) self-mapping counts, (ii) cross-mapping counts and (iii) divergence corrected counts (*yˆ_i_* Equation 1) from simulations based on the *in silico* and Ensembl genomes.

## Supplementary Material

Supplementary figures S1-S7 and Supplementary table S1 are given in the additional file.

## References

1. Wolfgang Enard, Philipp Khaitovich, Joachim Klose, Sebastian Zöllner, Florian Heissig, Patrick Giavalisco, Kay Nieselt-Struwe, Elaine Muchmore, Ajit Varki, Rivka Ravid, Gaby M Doxiadis, Ronald E Bontrop, and Svante Pääbo. Intra- and interspecific variation in primate gene expression patterns. Science, 296(5566):340–343, April 2002.

2. Julien Roux, Marta Rosikiewicz, and Marc Robinson-Rechavi. What to compare and how: Comparative transcriptomics for Evo-Devo. J. Exp. Zool. B Mol. Dev. Evol., 324(4):372–382, 2015.

3. Hunter B Fraser. Genome-wide approaches to the study of adaptive gene expression evolution. Bioessays, 33(6):469–477, 2011.

4. Anamaria Necsulea and Henrik Kaessmann. Evolutionary dynamics of coding and non-coding transcriptomes. Nat. Rev. Genet., 15(11):734–748, November 2014.

5. Athma A Pai, Jonathan K Pritchard, and Yoav Gilad. The genetic and mechanistic basis for variation in gene regulation. PLoS Genet., 11(1):e1004857, January 2015.

6. Armita Nourmohammad, Joachim Rambeau, Torsten Held, Viera Kovacova, Johannes Berg, and Michael Lässig. Adaptive evolution of gene expression in drosophila. Cell Rep., 20(6):1385–1395, August 2017.

7. Philipp Khaitovich, Wolfgang Enard, Michael Lachmann, and Svante Pääbo. Evolution of primate gene expression. Nat. Rev. Genet., 7(9):693–702, September 2006.

8. Philipp Khaitovich, Bjoern Muetzel, Xinwei She, Michael Lachmann, Ines Hellmann, Janko Dietzsch, Stephan Steigele, Hong-Hai Do, Gunter Weiss, Wolfgang Enard, Florian Heissig, Thomas Arendt, Kay Nieselt-Struwe, Evan E Eichler, and Svante Pääbo. Regional patterns of gene expression in human and chimpanzee brains. Genome Res., 14(8):1462–1473, August 2004.

9. Yoav Gilad, Scott A Rifkin, Paul Bertone, Mark Gerstein, and Kevin P White. Multi-species microarrays reveal the effect of sequence divergence on gene expression profiles. Genome Res., 15(5): 674–680, May 2005.

10. Scott A Rifkin, Junhyong Kim, and Kevin P White. Evolution of gene expression in the drosophila melanogaster subgroup. Nat. Genet., 33(2):138–144, February 2003.

11. Michael Dannemann, Anna Lorenc, Ines Hellmann, Philipp Khaitovich, and Michael Lachmann. The effects of probe binding affinity differences on gene expression measurements and how to deal with them. Bioinformatics, 25(21):2772–2779, November 2009.

12. Yu Zhang, David Sturgill, Michael Parisi, Sudhir Kumar, and Brian Oliver. Constraint and turnover in sex-biased gene expression in the genus drosophila. Nature, 450(7167):233–237, November 2007.

13. Ashlee M Benjamin, Marshall Nichols, Thomas W Burke, Geoffrey S Ginsburg, and Joseph E Lucas. Comparing reference-based RNA-Seq mapping methods for non-human primate data. BMC Genomics, 15(1):570, December 2014.

14. Ran Blekhman, John C Marioni, Paul Zumbo, Matthew Stephens, and Yoav Gilad. Sex-specific and lineage-specific alternative splicing in primates. Genome Res., 20(2):180–189, February 2010.

15. David Brawand, Magali Soumillon, Anamaria Necsulea, Philippe Julien, Gábor Csárdi, Patrick Harrigan, Manuela Weier, Angélica Liechti, Ayinuer Aximu-Petri, Martin Kircher, Frank W Albert, Ulrich Zeller, Philipp Khaitovich, Frank Grützner, Sven Bergmann, Rasmus Nielsen, Svante Pääbo, and Henrik Kaessmann. The evolution of gene expression levels in mammalian organs. Nature, 478(7369): 343–348, October 2011.

16. Ying Zhu, Mingfeng Li, André Mm Sousa, and Nenad Sestan. XSAnno: a framework for building ortholog models in cross-species transcriptome comparisons. BMC Genomics, 15:343, May 2014.

17. André M M Sousa, Ying Zhu, Mary Ann Raghanti, Robert R Kitchen, Marco Onorati, Andrew T N Tebbenkamp, Bernardo Stutz, Kyle A Meyer, Mingfeng Li, Yuka Imamura Kawasawa, Fuchen Liu, Raquel Garcia Perez, Marta Mele, Tiago Carvalho, Mario Skarica, Forrest O Gulden, Mihovil Pletikos, Akemi Shibata, Alexa R Stephenson, Melissa K Edler, John J Ely, John D Elsworth, Tamas L Horvath, Patrick R Hof, Thomas M Hyde, Joel E Kleinman, Daniel R Weinberger, Mark Reimers, Richard P Lifton, Shrikant M Mane, James P Noonan, Matthew W State, Ed S Lein, James A Knowles, Tomas Marques-Bonet, Chet C Sherwood, Mark B Gerstein, and Nenad Sestan. Molecular and cellular reorganization of neural circuits in the human lineage. Science, 358(6366):1027–1032, November 2017.

18. Stephanie Wunderlich, Martin Kircher, Beate Vieth, Alexandra Haase, Sylvia Merkert, Jennifer Beier, Gudrun Gohring, Silke Glage, Axel Schambach, Eliza C Curnow, Svante Pääbo, Ulrich Martin, and Wolfgang Enard. Primate iPS cells as tools for evolutionary analyses. Stem Cell Res., 12(3):622–629, May 2014.

19. Benedict Paten, Javier Herrero, Kathryn Beal, Stephen Fitzgerald, and Ewan Birney. Enredo and pecan: genome-wide mammalian consistency-based multiple alignment with paralogs. Genome Res., 18(11):1814–1828, November 2008.

20. Albert J Vilella, Jessica Severin, Abel Ureta-Vidal, Li Heng, Richard Durbin, and Ewan Birney. EnsemblCompara GeneTrees: Complete, duplication-aware phylogenetic trees in vertebrates. Genome Res., 19(2):327–335, February 2009.

21. Thomas Derrien, Jordi Estellé, Santiago Marco Sola, David G Knowles, Emanuele Raineri, Roderic Guigo, and Paolo Ribeca. Fast computation and applications of genome mappability. PLoS One, 7 (1):e30377, January 2012.

22. Robert C Edgar, George Asimenos, Serafim Batzoglou, and Arend Sidow. Evolver. Website http://www.drive5.com/evolver, 2009.

23. Hervé Philippe, Henner Brinkmann, Dennis V Lavrov, D Timothy J Littlewood, Michael Manuel, Gert Worheide, and Denis Baurain. Resolving difficult phylogenetic questions: why more sequences are not enough. PLoS Biol., 9(3):e1000602, March 2011.

24. Thasso Griebel, Benedikt Zacher, Paolo Ribeca, Emanuele Raineri, Vincent Lacroix, Roderic Guigó, and Michael Sammeth. Modelling and simulating generic RNA-Seq experiments with the flux simulator. Nucleic Acids Res., 40(20):10073–10083, November 2012.

25. Javier Herrero, Matthieu Muffato, Kathryn Beal, Stephen Fitzgerald, Leo Gordon, Miguel Pignatelli, Albert J Vilella, Stephen M J Searle, Ridwan Amode, Simon Brent, William Spooner, Eugene Kulesha, Andrew Yates, and Paul Flicek. Ensembl comparative genomics resources. Database, 2016, February 2016.

26. Alexander Dobin, Carrie A Davis, Felix Schlesinger, Jorg Drenkow, Chris Zaleski, Sonali Jha, Philippe Batut, Mark Chaisson, and Thomas R Gingeras. STAR: ultrafast universal RNA-seq aligner. Bioinformatics, 29(1):15–21, January 2013.

27. Giacomo Baruzzo, Katharina E Hayer, Eun Ji Kim, Barbara Di Camillo, Garret A FitzGerald, and Gregory R Grant. Simulation-based comprehensive benchmarking of RNA-seq aligners. Nat. Methods, December 2016.

28. Michael I Love, Wolfgang Huber, and Simon Anders. Moderated estimation of fold change and dispersion for RNA-seq data with DESeq2. Genome Biol., 15(12):550, 2014.

29. SEQC/MAQC-III Consortium. A comprehensive assessment of RNA-seq accuracy, reproducibility and information content by the sequencing quality control consortium. Nat. Biotechnol., 32(9):903–914, September 2014.

30. Matthew E Ritchie, Belinda Phipson, Di Wu, Yifang Hu, Charity W Law, Wei Shi, and Gordon K Smyth. limma powers differential expression analyses for RNA-sequencing and microarray studies. Nucleic Acids Res., 43(7):e47, April 2015.

31. Erica V Todd, Michael A Black, and Neil J Gemmell. The power and promise of RNA-seq in ecology and evolution. Mol. Ecol., 25(6):1224–1241, March 2016.

32. Swati Parekh, Christoph Ziegenhain, Beate Vieth, Wolfgang Enard, and Ines Hellmann. The impact of amplification on differential expression analyses by RNA-seq. Sci. Rep., 6:25533, May 2016.

33. W James Kent, Charles W Sugnet, Terrence S Furey, Krishna M Roskin, Tom H Pringle, Alan M Zahler, and David Haussler. The human genome browser at UCSC. Genome Res., 12(6):996–1006, June 2002.

34. T Hubbard, D Barker, E Birney, G Cameron, Y Chen, L Clark, T Cox, J Cuff, V Curwen, T Down, R Durbin, E Eyras, J Gilbert, M Hammond, L Huminiecki, A Kasprzyk, H Lehvaslaiho, P Lijnzaad, C Melsopp, E Mongin, R Pettett, M Pocock, S Potter, A Rust, E Schmidt, S Searle, G Slater, J Smith, W Spooner, A Stabenau, J Stalker, E Stupka, A Ureta-Vidal, I Vastrik, and M Clamp. The ensembl genome database project. Nucleic Acids Res., 30(1):38–41, January 2002.

35. Beate Vieth, Christoph Ziegenhain, Swati Parekh, Wolfgang Enard, and Ines Hellmann. powsimr: Power analysis for bulk and single cell RNA-seq experiments. Bioinformatics, July 2017.

36. Robert M Kuhn, David Haussler, and W James Kent. The UCSC genome browser and associated tools. Brief. Bioinform., 14(2):144–161, March 2013.

37. Yang Liao, Gordon K Smyth, and Wei Shi. featurecounts: an efficient general purpose program for assigning sequence reads to genomic features. Bioinformatics, 30(7):923–930, April 2014.

38. Mark A DePristo, Eric Banks, Ryan Poplin, Kiran V Garimella, Jared R Maguire, Christopher Hartl, Anthony A Philippakis, Guillermo del Angel, Manuel A Rivas, Matt Hanna, Aaron McKenna, Tim J Fennell, Andrew M Kernytsky, Andrey Y Sivachenko, Kristian Cibulskis, Stacey B Gabriel, David Altshuler, and Mark J Daly. A framework for variation discovery and genotyping using next-generation DNA sequencing data. Nat. Genet., 43(5):491–498, May 2011.

39. M Kimura. A simple method for estimating evolutionary rates of base substitutions through comparative studies of nucleotide sequences. J. Mol. Evol., 16(2):111–120, December 1980.

40. Yoav Benjamini and Daniel Yekutieli. The control of the false discovery rate in multiple testing under dependency. Ann. Stat., 29(4):1165–1188, 2001.

